# Stress-adapted endophytic *Pseudomonas* strains isolated from unmanaged lupin promote potato growth and suppress verticillium wilt through complementary biocontrol and plant growth-promoting mechanisms

**DOI:** 10.64898/2026.11.06.764942

**Authors:** Numan Ibne Asad, Lia Watson, Anab Mujtaba, Ievgen Lebedenko, Selvakumari Arunachalam, Hamid Reza Eisvand, Haleema Tariq, Donald L. Smith

## Abstract

Verticillium wilt, primarily caused by *Verticillium dahliae*, is among the most destructive soil-borne diseases impacting global potato production. Fungal propagules remain viable in soil for extended periods, and conventional control measures often show limited efficacy. These challenges underscore the necessity for sustainable disease-management strategies. We evaluated two stress-adapted endophytic *Pseudomonas* strains, B18 and B14, isolated from unmanaged wild *Lupinus* roots in Québec, Canada, for their plant growth–promoting and biocontrol potential against Verticillium wilt in potato.

Both isolates exhibited multiple plant-beneficial traits. B18 demonstrated strong phosphate solubilization, indole-3-acetic acid production, putative nitrogen fixation-related activity, siderophore production, and hydrogen cyanide production. These traits correlated with enhanced plant growth–promoting effects, including stimulation of seed germination and radicle growth. In contrast, B14 primarily exhibited antagonistic activity against *Verticillium* species through similar plant-beneficial mechanisms, indicating complementary activities between the strains. The cell-free supernatants of B18 further supported its biostimulant activity, suggesting a significant role for its extracellular metabolites.

Greenhouse experiments using the *Verticillium*-susceptible potato cultivar Russet Burbank, inoculated with either strain, showed enhanced plant growth and reduced disease severity compared with pathogen-only controls. Under pathogen challenge, plants treated with plant growth–promoting rhizobacteria (PGPR) exhibited up to 62% greater shoot height and approximately double the leaf production. Additionally, these plants produced 4.2 times more tubers than pathogen-only plants. Disease progression, measured as the area under the disease progress curve (AUDPC), was reduced by 84–92% following bacterial inoculation. While individual strains effectively suppressed disease, the combined B18 plus B14 strains resulted in the most substantial improvements in growth, biomass, root development, and tuber production.

These findings demonstrate that stress-adapted endophytic *Pseudomonas* strains from wild *Lupinus* can contribute to sustainable potato production. The B18 plus B14 consortium, when combined with two-strain inoculation, demonstrated the most significant effects. These findings emphasize the potential of functionally complementary microbial inoculants for the integrated management of *Verticillium* wilt and other persistent soil-borne diseases in potatoes.

## 1. Introduction

Potato (*Solanum tuberosum* L.) ranks among the most globally important food crops due to its high nutritional value, adaptability, and economic significance. Despite its importance for food security, potato production is continually threatened by soil-borne pathogens that reduce both yield and tuber quality. *Verticillium* species, in particular, are responsible for Verticillium wilt, a destructive vascular disease prevalent in potato-growing regions worldwide. Infections by *Verticillium dahliae*, *V. longisporum*, and related species result in chlorosis, wilting, vascular discoloration, diminished tuber yield, and substantial economic losses, especially in intensive production systems (Pegg and Brady, 2002; Klosterman et al., 2009). The persistence of *Verticillium* microsclerotia in soil and the broad host range of these pathogens present significant challenges for effective disease management.

Current management strategies for Verticillium wilt primarily include crop rotation, the use of resistant cultivars, soil fumigation, and direct application of chemical fungicides. These methods, however, frequently yield inconsistent results, can be economically burdensome, and may contribute to environmental concerns due to intensive chemical use (Fradin and Thomma, 2006; Depotter et al., 2016). As a result, there is increasing interest in sustainable biological alternatives that support plant health and suppress pathogen development.

Plant growth-promoting rhizobacteria (PGPR) have emerged as promising agents for sustainable crop production and disease management. These beneficial microorganisms enhance plant performance through mechanisms such as nutrient mobilization, phytohormone production, biological nitrogen fixation, siderophore generation, pathogen suppression, and stress alleviation (Lugtenberg and Kamilova, 2009; Glick, 2012). Within the PGPR group, *Pseudomonas* species are notable for their metabolic versatility, environmental adaptability, efficient root colonization, and capacity to suppress a wide range of plant pathogens while promoting growth (Haas and Défago, 2005; Weller, 2007). Additionally, many *Pseudomonas* species produce diverse bioactive metabolites that inhibit pathogens and enhance plant performance, making them valuable for agricultural biotechnologies (Mercado-Blanco and Bakker, 2007; Pieterse et al., 2014).

Recent studies increasingly identify plant endophytes as reservoirs of agriculturally beneficial microorganisms. Unlike free-living rhizosphere bacteria, endophytes establish close associations within plant tissues, where they can directly enhance plant nutrition, stress tolerance, and disease resistance (Compant et al., 2010; Hardoim et al., 2015). Wild and unmanaged plant populations in challenging environments frequently harbor microbiomes with traits associated with adaptation and plant fitness. These environments represent valuable yet underexplored sources of microbial resources for sustainable agriculture (Rodríguez et al., 2009; Trivedi et al., 2020).

Accumulating evidence from previous literature indicates that microbial consortia frequently outperform single-strain inoculants, as diverse microorganisms can provide complementary ecological functions that enhance plant performance (Bashan et al., 2014; Niu et al., 2017; Berg et al., 2021). Nevertheless, it remains uncertain whether functionally complementary endophytic *Pseudomonas* strains from stress-adapted wild hosts can simultaneously promote potato growth and suppress Verticillium wilt. Furthermore, the potential of these isolates to serve as sources of biologically active extracellular metabolites is not well understood. This study investigates the effects of such *Pseudomonas* consortia on potato performance and disease suppression, with a specific focus on their dual functional potential.

In this study, we isolated endophytic *Pseudomonas* strains from the roots of unmanaged wild lupin plants growing in challenging environments and evaluated their growth-promoting and biocontrol potential. We hypothesized that stress-adapted isolates would possess traits that enhance plant growth and suppress Verticillium wilt, and that combining strains with complementary functions would be more effective than using single strains. To test these hypotheses, we (i) characterized growth-promoting traits and antifungal activity, (ii) evaluated the biological activity of cell-free supernatants, (iii) assessed effects on seed germination and seedling development, and (iv) tested the individual and combined ability of these strains to promote potato growth and reduce *Verticillium wilt* under greenhouse conditions.

## 2. Materials and methods

### 2.1 Plant sampling and selection of candidate endophytes

Healthy wild *Lupinus* plants were collected from the lakefront area of the Macdonald Campus of McGill University, Ste. Anne de Bellevue, Québec, Canada, in December 2024. Sampling focused on plants from exposed field sites that persisted during seasonal low temperatures (below 0 °C) and on nutrient-limited soils. Plants thriving under such environmental stress frequently harbor endophytic microorganisms with enhanced adaptive and plant-beneficial traits. Therefore, the sustained vigor of *Lupinus* spp. under these conditions suggests that their root microbiome represents a promising source of stress-adapted plant growth–promoting rhizobacteria (PGPR) for subsequent greenhouse and field applications.

### 2.2 Isolation and preservation of endophytic bacteria

Root samples were thoroughly washed under running tap water to remove adhering soil particles and surface-sterilized by sequential immersion in 75% ethanol for 10 min, 3% sodium hypochlorite for 3–5 min, and sterile distilled water six times. To verify successful removal of epiphytic microorganisms, aliquots of the final rinse water were plated onto King’s B medium (King et al., 1954), tryptic soy agar (TSA), and nutrient agar (NA), and incubated to confirm the absence of microbial growth. Sterilized root tissues were aseptically macerated in sterile phosphate-buffered saline (PBS), serially diluted (10^-1^ to 10^-3^), and spread onto the same media. Plates were incubated at 28°C for 3–5 d. Colonies differing in morphology, pigmentation, size, or texture were purified by repeated (3 times) streaking to obtain axenic cultures.

Pure isolates were subsequently grown in nutrient broth at 28°C with shaking (150 rpm) for 24–48 h to reach a cell density of approximately 10^8^ to 10^9^ CFU mL^-1^, estimated by OD₆₀₀ measurements. Broth cultures were preserved in 50% glycerol at −80°C for long-term storage. To prepare bacterial inoculum for downstream analyses, cells were harvested by centrifugation (6000 × *g*, 10 min), washed twice, and resuspended in sterile 10 mM MgSO₄ solution. Cell density was then adjusted to approximately 10^8^ CFU mL^-1^ using OD₆₀₀ measurements. The MgSO₄ solution was used as a low-nutrient carrier to maintain osmotic stability and bacterial cell viability during inoculation procedures.

### 2.5 Screening of biocontrol isolates by *in vitro* antagonistic activity against Verticillium spp

A total of 24 distinct bacterial isolates recovered from lupin root endophytes were initially screened for antagonistic activity against *Verticillium* spp. using a dual-culture inhibition assay on potato dextrose agar (PDA), a widely used preliminary method for identifying microbial candidates with antifungal potential (Dennis and Webster, 1971; Weller, 2007). Sterile 90-mm Petri dishes containing PDA were prepared, and a sterile filter paper disc (4.5 cm diameter) was placed on the agar surface to standardize inoculation positions and ensure uniform spacing among treatments.

A 5-mm mycelial plug excised from the actively growing margin of the fungal colony was positioned at the center of each plate. Four equidistant inoculation points, approximately 2 cm from the fungal plug, were marked around the center. At each point, 10 µL of bacterial suspension (approximately 1 × 10^8^ CFU mL^-1^) was applied. Control plates received the fungal plug alone without bacterial inoculation. Plates were incubated at 28°C for 5 to 7 days and observed regularly for fungal growth restriction, formation of inhibition zones, and visible changes in colony morphology (Fig. S1).

At the end of the incubation period, fungal colony diameter was measured in millimetres using two perpendicular axes across the colony through the center of the inoculation point, and the mean value was used for analysis, as commonly applied in mycelial growth inhibition assays (Pandey *et al*., 1982; Dennis and Webster, 1971). Percentage inhibition of radial growth was calculated according to the formula:

Inhibition (%) = [(fungal colony diameter in control (C) − fungal colony diameter in dual culture (T) / fungal colony diameter in control (C)] × 100

Where **C** is the mean colony diameter of the fungal control plate, and **T** is the mean colony diameter of the fungus in dual culture treatment (Vincent, 1947).

Each treatment was performed in triplicate using three independent biological replicates. Isolates were comparatively ranked based on qualitative observations, such as clear suppression zones, restricted mycelial expansion, and altered fungal growth patterns, as well as quantitative inhibition percentages.

Isolates that consistently exhibited the strongest and most reproducible antagonistic activity against *Verticillium spp*. were selected for further investigation. These isolates underwent functional characterization of plant growth–promoting traits, seed bioassays, molecular identification, and greenhouse evaluation of biological control efficacy in potato.

### 2.4 Functional screening for plant growth–promoting traits

The isolates identified in the dual culture assay as having the most antagonistic potential were further evaluated for multiple plant growth–promoting (PGP) traits using established standard biochemical assays. These assays were selected to assess functional mechanisms associated with nutrient acquisition, phytohormone production, micronutrient mobilization, and pathogen suppression. Appropriate positive and negative controls were included in each assay to ensure reliability and consistency of interpretation (Fig. S2).

Indole-3-acetic acid (IAA) production was assessed qualitatively and quantitatively following the method of Gordon and Weber (1951) with modifications. Isolates were grown in tryptic soy broth (TSB) supplemented with L-tryptophan (0.5 to 1 g L^-1^) at 28°C for 48 h. Cultures were then centrifuged, and 1 mL of supernatant was mixed with 2 mL of Salkowski reagent. Development of pink coloration indicated IAA production. Quantitative estimation was performed by measuring absorbance at 530 nm and calculating IAA concentration from a standard curve (Patten and Glick, 2002).

Phosphate solubilization ability was evaluated on Pikovskaya’s agar containing insoluble tricalcium phosphate (Pikovskaya, 1948). Bacterial cultures were spot-inoculated and incubated at 28°C for 5 to 7 days. The formation of clear halo zones around colonies indicated the solubilization of inorganic phosphate. Solubilization efficiency was expressed as the phosphate solubilization index (PSI), calculated as the ratio of total diameter (colony + halo zone) to colony diameter.

Nitrogen fixation-associated activity was evaluated qualitatively using a nitrogen-free bromothymol blue (NFb) semi-solid medium, as described by Döbereiner (1995). Isolates were inoculated into NFb medium and incubated at 28°C for 3 to 5 days. Development of bacterial growth, accompanied by a colour change from green to blue, was interpreted as evidence of growth under nitrogen-limited conditions and of ammonia production associated with potential nitrogen-fixing capability.

Because this assay provides only preliminary qualitative evidence, results are reported as nitrogen fixation-associated activity rather than direct confirmation of biological nitrogen fixation. But definitive confirmation of nitrogen fixation might require further assay like acetylene reduction, ^15^N isotope incorporation experiments, or molecular characterization of nitrogenase-related genes.

Hydrogen cyanide (HCN) production, an important trait linked to suppression of soil-borne pathogens, was assessed following Bakker and Schippers (1987). Isolates were grown on TSB agar supplemented with glycine, and filter paper impregnated with picric acid solution was placed in the lid of each plate. A colour change from yellow to brown after 2 to 4 days of incubation indicated HCN production.

Siderophore production was evaluated using Chrome Azurol S (CAS) agar, as described by Schwyn and Neilands (1987). The formation of orange halos surrounding colonies indicated the secretion of iron-chelating siderophores.

These assays aimed to provide preliminary functional characterization of the isolates and to identify traits commonly associated with plant growth promotion and biological control. The assays were designed to demonstrate functional potential, not to quantify the contribution of individual mechanisms under greenhouse conditions.

### 2.5 Seed germination bioassays

Tomato seeds served as a rapid and standardized model system to evaluate the effects of bacterial inoculation on seed germination and early seedling development. Both tomato and potato are members of the *Solanaceae* family and share similar physiological characteristics during early development. Unlike potato, which is propagated from nutrient-rich tubers that may obscure microbial effects during establishment, tomato seeds offer a sensitive and uniform system for detecting growth-promoting responses. Additionally, tomato seeds germinate rapidly and synchronously under controlled conditions, making them suitable for preliminary screening of plant growth-promoting microorganisms prior to validation in the target crop.

Surface-sterilized seeds were placed on sterile filter paper in Petri dishes and treated with bacterial suspensions adjusted to approximately 10^5^, 10^6^, or 10^7^ CFU mL^-1^. Sterile 10 mM MgSO_4_ served as the carrier control. Each treatment consisted of four biological replicates, with 25 seeds per Petri dish. Each Petri dish was considered a single experimental unit.

Seeds were incubated under controlled growth conditions, and germination was recorded daily for three days. A seed was considered germinated when visible radicle emergence occurred. Germination percentage was calculated on a per-plate basis. Germination speed was assessed using Maguire’s Germination Index (Maguire, 1962):

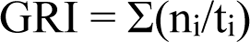

where n_i_ represents the number of newly germinated seeds at time tᵢ.

### 2.6 Cell-free supernatant assays

To determine whether extracellular metabolites contributed to the observed plant responses independently of bacterial colonization, cell-free supernatants (CFS) were prepared from liquid cultures of the selected isolates. Cultures were grown at 28°C for 24 to 48 hours, centrifuged at 10,000 × g for 10 min, and filtered through sterile 0.22 μm membranes to remove bacterial cells.

Supernatants were diluted with sterile distilled water to final concentrations of 2.5, 5, and 10% (v/v). Two controls were included: (i) sterile distilled water and (ii) uninoculated sterile culture medium processed identically to the CFS preparations. These controls enabled discrimination between the effects of bacterial metabolites and residual medium components.

### 2.8 Molecular identification and phylogenetic analysis

Forward and reverse partial 16S rRNA gene sequences of the selected lupin endophytic isolates were obtained by Sanger sequencing at Genome Québec following submission of pure bacterial culture samples. Raw reads were quality filtered, trimmed, and assembled into consensus sequences. Closely related type and reference sequences were retrieved from NCBI for comparative analysis.

Multiple sequence alignment was performed using the R package msa (Bodenhofer et al., 2015). Phylogenetic relationships were inferred using maximum likelihood in phangorn (Schliep, 2011) under the GTR+G+I substitution model. Branch support was assessed using 200 bootstrap replicates. Trees were rooted using *Azotobacter vinelandii* and visualized using ggtree (Yu et al., 2017) and iTOLV6 (Letunic and Bork 2024).

### 2.9 Greenhouse evaluation of biocontrol efficacy against *Verticillium dahliae*

#### 2.9.1 Experimental design and treatments

Greenhouse experiments were conducted using Agromix substrate in a randomized complete block design (RCBD) to evaluate the preventive biocontrol efficacy of selected plant growth–promoting rhizobacteria (PGPR) against *Verticillium dahliae* in potato. The experiment consisted of eight treatment combinations, each with eight biological replicates, resulting in a total of 64 experimental units (pots). Experimental units were arranged into blocks, with each block containing all treatment combinations randomly assigned using a random number generator to minimize positional effects and account for environmental variability within the greenhouse.

The potato cultivar Russet Burbank, known to be susceptible to *Verticillium* wilt and commonly used in disease screening studies, was selected to ensure consistent disease development and maximize sensitivity for detecting treatment effects.

Based on the preliminary screening results, strain B18 was designated PGPR A and strain B14 was designated PGPR B. The greenhouse experiment followed a 4 × 2 factorial arrangement, consisting of four PGPR treatments [no PGPR (control), PGPR A (B18), PGPR B (B14), and PGPR A + B (B18 + B14)] and two pathogen levels [absence or presence of *Verticillium dahliae*]. The combination of these factors generated eight treatment combinations: (i) healthy control, (ii) PGPR A only, (iii) PGPR B only, (iv) PGPR A + B only, (v) pathogen only, (vi) PGPR A + pathogen, (vii) PGPR B + pathogen, and (viii) PGPR A + B + pathogen. The healthy control received a sterile 10 mM MgSO_4_ carrier solution supplemented with 0.25% carboxymethyl cellulose (CMC) and no bacterial inoculum, whereas pathogen-free PGPR treatments received only a bacterial inoculum. Pathogen treatments received *V. dahliae* inoculation either alone or in combination with the corresponding PGPR treatment.

This treatment structure allowed for the separation of direct plant growth–promoting effects under healthy conditions from disease-protective effects under pathogen pressure. It also enabled evaluation of potential additive or synergistic effects resulting from combined inoculation.

#### 2.9.2 Preparation and application of bacterial inoculum

Healthy potato tubers were selected, cut into uniform cubes (∼ 5g), surface-sterilized, and air-dried prior to treatment to minimize background microbial interference. Bacterial inoculum was prepared and adjusted to a cell concentration of approximately 10⁸ CFU mL⁻¹ in sterile 10 mM MgSO_4_ buffer, which served as a low-nutrient carrier to maintain osmotic stability and cell viability without introducing excess nutrients (Bashan *et al*., 2014). The PGPR A+B consortium was prepared by cultivating each strain (B18 and B14) separately, standardizing the resulting cell suspensions to an equivalent cell density, and subsequently combining equal volumes of the two suspensions immediately prior to inoculation. This method produced a dual-strain consortium of B18 and B14 at a 1:1 ratio, aligning with established PGPR consortium inoculation protocols in which individually prepared strains are standardized and combined before application (Hashmi et al., 2019; Khan et al., 2022; Petrillo et al., 2022).

Tubers assigned to bacterial treatments were inoculated by immersion in the respective suspension for 1 h. The inoculum suspension contained 0.25% carboxymethyl cellulose (CMC) as a sticker to improve bacterial adhesion and facilitate early colonization of the tuber surface.

To enhance rhizosphere establishment, a second application of bacterial inoculum was delivered as a soil drench on Day +3 after planting (10 mL pot^-1^ at approximately 10^8^ CFU mL^-1^).

#### 2.9.3 Plant nutrition and irrigation management

A minimal hydroponic nutrient input consisting of stock solutions A and B was applied during the vegetative stage at approximately 0.0000625% (v/v; ∼0.6 ppm) to maintain baseline plant nutrition while minimizing interference with microbial interactions. Solution A primarily contained calcium nitrate and iron chelate, whereas Solution B contained phosphate, magnesium sulphate, potassium salts, and trace micronutrients (B, Mn, Zn, Cu, and Mo).

Prior to pathogen inoculation and throughout the experiment, all pots received autoclaved distilled water as needed to maintain consistent soil moisture across treatments.

#### 2.9.4 Preparation and application of pathogen inoculum

The pathogen inoculum consisted of *V. dahliae* conidia harvested from potato dextrose agar (PDA) cultures using sterile distilled water supplemented with Tween-20 to facilitate spore release. The suspension was filtered to remove mycelial fragments and quantified using a hemocytometer under light microscopy (Lievens *et al*., 2006). Spore concentration was adjusted to approximately 10⁶ conidia mL⁻¹.

To improve infection consistency and better simulate natural inoculum pressure, actively growing fungal mycelial plugs (5 mm diameter) were also introduced into the soil near the root zone. Pathogen inoculation was performed approximately one week after plant emergence, when seedlings were 1 to 2 cm tall and had reached a uniform, susceptible growth stage. This time point was considered the baseline for Verticillium infection assessment. Inoculum was applied as a soil drench (10 mL pot^-1^) to allow direct contact between fungal propagules and the rhizosphere.

#### 2.9.5 Greenhouse Conditions

Plants were maintained under controlled greenhouse conditions of 22/17°C Day/night temperature, a 16 hours photoperiod, and 60–70% relative humidity. Irrigation was adjusted as required to maintain uniform moisture among all treatments.

#### 2.9.6 Plant growth and disease assessments

Plant emergence was recorded during establishment and expressed as emergence percentage per treatment using the formula:

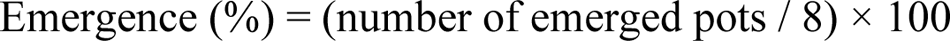

Because each planted tuber piece was capable of producing multiple shoots, the total number of shoots emerging per seed tuber was also recorded as an indicator of sprouting vigour and stem multiplication.

Vegetative growth parameters, including shoot height (cm) and total leaf number per experimental unit, were measured at emergence (0 dpi) and subsequently at 7, 14, 21, and 28 days post-inoculation (dpi). Shoot height was measured from the substrate surface to the apex of the tallest stem using a ruler, while leaf number included all fully expanded leaves arising from each planted tuber.

Leaf greenness, used as an indirect indicator of chlorophyll status and physiological performance, was measured non-destructively using a SPAD chlorophyll meter (SPAD-502, Konica Minolta, Japan or equivalent) on the youngest fully expanded leaves. Total leaf area was determined at harvest using a portable leaf area meter (CI-202/CI-203 series, CID Bio-Science, USA), which measures projected leaf surface area (cm^2^) photometrically.

#### 2.9.7 Disease severity and final harvest measurements

Disease severity was visually assessed at 7, 14, 21, and 28 days post-pathogen inoculation (dpi) using a standardized 0–5 wilt severity scale based on foliar chlorosis, wilting, necrosis, and canopy collapse, where: 0 = asymptomatic healthy plant; 1 = 1–25% foliage affected; 2 = 26–50% affected with slight stunting; 3 = 51–75% affected with moderate wilting and necrosis; 4 = >75% affected with severe wilt and chlorosis; and 5 = plant collapse or near death. To ensure scoring consistency, all assessments were performed by the same evaluator.

Disease progression over time was summarized as the Area Under the Disease Progress Curve (AUDPC), as described by Shaner and Finney (1977). At final harvest, plants were carefully removed from the substrate and separated into shoots, roots, and tubers. Fresh biomass was recorded immediately, then oven-dried to a constant weight for dry biomass determination. Flowering incidence was determined by scoring each pot as flowering (1) or non-flowering (0) based on the presence of at least one visible inflorescence. Flowering incidence was expressed as the percentage of pots that flowered within each treatment group. Tuber bulking response was expressed as the number of tubers produced per seed tuber planted. Root system architecture, including total root length, root surface area, root volume, and average root diameter, was quantified using a flatbed scanner coupled with WinRHIZO image analysis software (Regent Instruments Inc., Canada).

Collectively, these measurements quantified the effects of PGPR treatments on plant establishment, growth promotion, physiological status, yield formation, root development, and suppression of *Verticillium* wilt.

### 2.10 Statistical Analysis

All statistical analyses were performed using R version 4.3.0 (R Core Team, 2023). Data were visually inspected and evaluated for normality and homogeneity of variance using residual diagnostic plots, Shapiro–Wilk tests, and Levene’s tests. When necessary, proportional data were arcsine square-root transformed prior to analysis to satisfy model assumptions.

Greenhouse experiments were analyzed using analysis of variance (ANOVA) within a randomized complete block design (RCBD), with treatment considered a fixed effect and block included as a random factor. Variables measured at a single time point were analyzed using one-way ANOVA. For variables measured repeatedly through time, observations were analyzed separately at each sampling date to facilitate biological interpretation of treatment responses. Seed germination and cell-free supernatant (CFS) experiments were analyzed using one-way ANOVA for each strain and treatment group.

When significant treatment effects were detected (*P* < 0.05), pairwise comparisons among means were performed using Tukey’s honestly significant difference (HSD) test at an experiment-wise significance level of α = 0.05. Adjusted *P*-values generated by Tukey’s multiple-comparison procedure were used to control the family-wise error rate associated with multiple testing. Treatment means sharing the same letter were not significantly different at *P* ≥ 0.05, whereas means with different letters were considered significantly different at *P* < 0.05. Estimated marginal means (EMMeans) and corresponding 95% confidence intervals were calculated using the emmeans package. Compact letter displays (CLD) were generated to summarize multiple-comparison results.

All figures were prepared using ggplot2 (Wickham, 2016). Results are presented as means ± standard error (SE) or means with 95% confidence intervals (CI), as indicated in the respective figures and tables. Statistical significance was declared at *P* < 0.05.

The R scripts used for data processing, statistical analyses, and figure generation are publicly available on GitHub (https://github.com/numanibneasad/potato-pgpr-b18-b14), whereas the experimental datasets, supplementary statistical summaries, and supporting files associated with this study are available through Zenodo (https://zenodo.org/records/20542212?preview=1).

## Results

### 3.1 Isolation and selection of candidate endophytic bacteria

Endophytic bacteria were isolated from surface-sterilized root tissues of wild *Lupinus* plants collected under cold-stress field conditions. The absence of microbial growth from the final rinse water plated on both selective and non-selective media confirmed effective surface sterilization and indicated the internal origin of the recovered isolates. Initial antagonistic screening against *Verticillium* spp. identified four candidate isolates with measurable inhibitory activity. Among these, isolates B14 and B18 consistently demonstrated the most promising characteristics and were selected for further investigation. Both isolates maintained stable growth during repeated subculturing and exhibited strong plant growth–promoting and biocontrol potential, justifying their selection for detailed characterization and greenhouse evaluation.

### 3.3 *In vitro* suppression of *Verticillium* spp

Building on the observed pathogen suppression traits, both isolates were next evaluated for their ability to inhibit mycelial growth of *Verticillium* species (Table 2).

Both isolates significantly inhibited mycelial growth of all tested *Verticillium* species (Table 2). Against *V. dahliae,* B14 caused 48.2% inhibition, compared with 37.4% for B18. Similar trends were observed for *V. longisporum*, where B14 and B18 reduced fungal growth by 55.8% and 46.0%, respectively.

Against *V. albo-atrum*, B14 again produced greater suppression (43.2%) than B18 (32.9%). Across all pathogens tested, B14 consistently outperformed B18 in direct antifungal activity. These data indicate that both strains possess broad-spectrum antagonistic potential, with B14 showing stronger direct suppression of *Verticillium* spp. under *in vitro* conditions. This led to further study of their effects on plant physiological processes.

### 3.2 Plant growth–promoting functional traits

The two isolates, B14 and B18, were further tested for plant-beneficial traits, with each displaying distinct functional profiles (Table 1). Strains B18 and B14 significantly solubilized inorganic phosphate, with phosphate solubilization index (PSI) values of 2.30 and 2.10, respectively, compared with the non-inoculated control. No significant difference was detected between the two strains.

**Table 1:** In *vitro* antagonistic activity of endophytic *Pseudomonas* strains B18 and B14 against three *Verticillium* pathogens in dual-culture assays. Values represent mean fungal radial growth (mm) from three biological replicates measured after 2 weeks of incubation. Different letters indicate significant differences within each pathogen according to Tukey’s HSD test (*P* < 0.05). Dash (—) indicates reference treatment or not applicable.

| **Pathogen** | **Treat ment** | **Radial growth (mm)** | **Control growth (mm)** | **Growth reduction (mm)** | **Inhibition (%)** | **Significance vs control** | ***p*-value** | **Tukey group** |
|---|---|---|---|---|---|---|---|---|
| *V. dahliae* | B18 | 23.74 ± SE | 37.95 | 14.21 | 37.44 | ** | 0.0034 | b |
| *V. dahliae* | B14 | 19.66 ± SE | 37.95 | 18.29 | 48.19 | *** | 0.0004 | b |
| *V. dahliae* | Control | 37.95 ± SE | 37.95 | 0.00 | 0.00 | — | — | a |
| *V. longisporum* | B18 | 22.00 ± SE | 40.74 | 18.74 | 46.00 | *** | 0.0005 | b |
| *V. longisporum* | B14 | 18.00 ± SE | 40.74 | 22.74 | 55.82 | *** | <0.0001 | b |
| *V. longisporum* | Control | 40.74 ± SE | 40.74 | 0.00 | 0.00 | — | — | a |
| *V. albo-atrum* | B18 | 26.00 ± SE | 38.74 | 12.74 | 32.89 | ** | 0.0014 | b |
| *V. albo-atrum* | B14 | 22.00 ± SE | 38.74 | 16.74 | 43.21 | *** | 0.0001 | b |
| *V. albo-atrum* | Control | 38.74 ± SE | 38.74 | 0.00 | 0.00 | — | — | a |

**Table 2.** Screening of plant growth–promoting (PGP) traits of bacterial isolates B18 and B14. Key plant growth–promoting traits of B18 and B14 were assessed both qualitatively and quantitatively. Indole-3-acetic acid (IAA) production was quantified spectrophotometrically (µg mL⁻¹). Phosphate solubilization efficiency is presented as the phosphate solubilization index (PSI), based on halo development on Pikovskaya’s agar. Nitrogen fixation, hydrogen cyanide (HCN) production, and siderophore production were evaluated qualitatively and are shown as either present (+) or absent (−). All measurements were under controlled laboratory conditions. Values represent mean observations across (n=3) biological replicates.

| Isolate | PSI | CLD | IAA (µg mL<sup>-1</sup>) | CLD | Nitrogen Fixation | HCN | CAS |
|---|---|---|---|---|---|---|---|
| Strain B14 | 2.10 | a | 0.18 | b | +++ | ++ | ++ |
| Strain B18 | 2.30 | a | 0.25 | a | +++ | +++ | +++ |
| Control | 1.00 | b | 0.06 | c | – | – | – |

Quantitative evaluation revealed that IAA production differed significantly among treatments. Strain B18 produced the highest IAA concentration (0.25 µg mL⁻¹), followed by B14 (0.18 µg mL⁻¹), whereas the control showed only negligible background levels (0.06 µg mL^-1^). Both isolates showed strong nitrogen fixation (+++). Strain B18 showed higher HCN and siderophore activity (+++) than B14 (++), suggesting greater potential for pathogen suppression and micronutrient mobilization.

Overall, the findings demonstrate that B14 and B18 possess complementary functional characteristics. B14 exhibits stronger antifungal activity, whereas B18 demonstrates greater plant growth–promoting capacity. Therefore, these two isolates were selected for subsequent seed germination, cell-free supernatant, and greenhouse experiments.

### 3.2 Seed germination and seed vigour responses to bacterial inoculation

Seed germination responded strongly to bacterial inoculation in a strain-dependent manner (Fig. 1). Overall, strain B18 exhibited substantially greater germination-promoting activity than strain B14, enhancing both germination speed and final germination percentage across inoculum concentrations.

**Figure 1.**
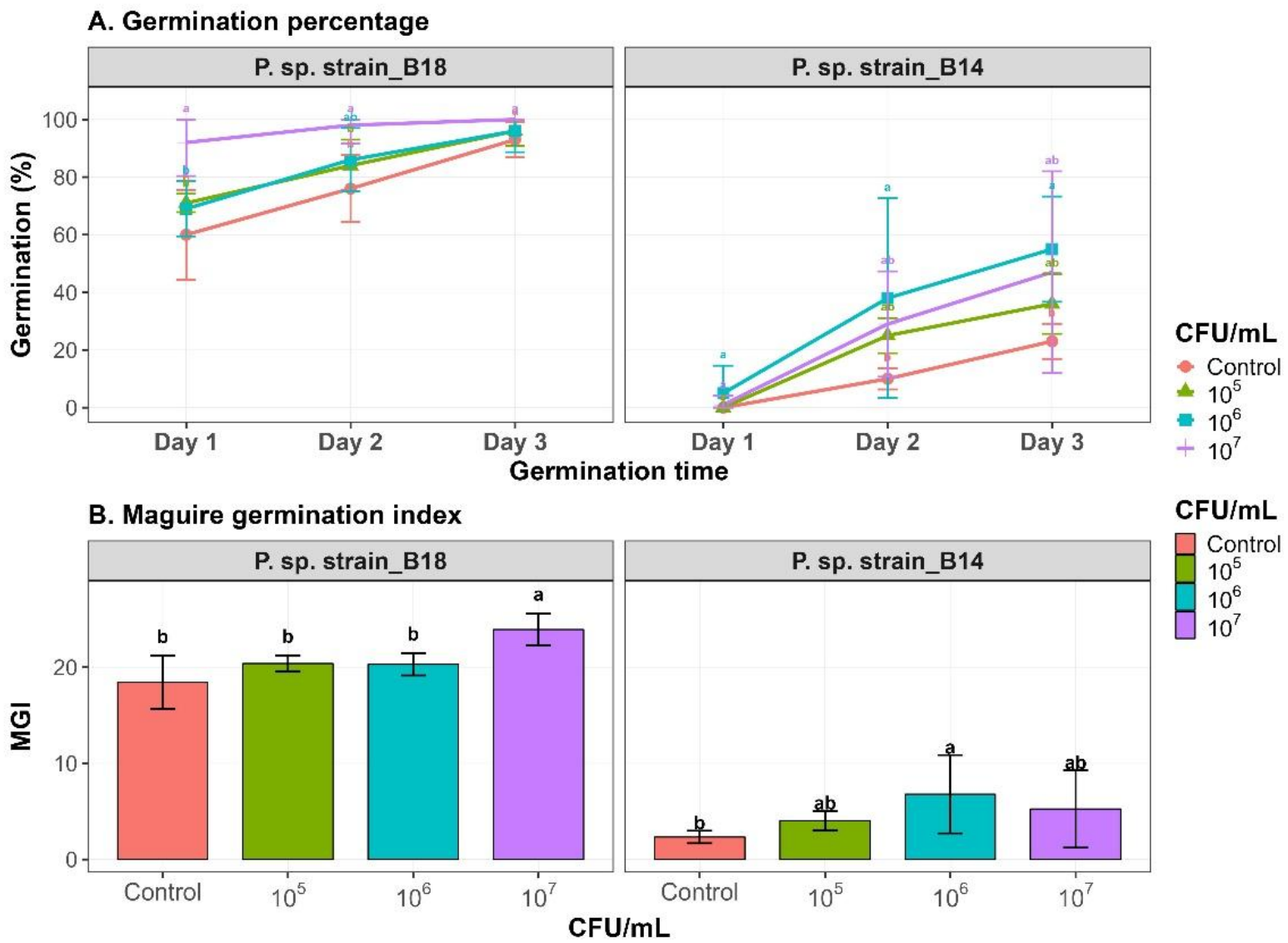
Effect of endophytic *Pseudomonas* strains on tomato seed germination and germination speed. Cumulative germination percentage and Maguire germination index (MGI) of tomato seeds treated with *P. sp.* strain_B18 and *P. sp.* strain_B14 at Control, 10^5^, 10^6^, and 10^7^ CFU mL^-1^. The control treatment contained sterile 10 mM MgSO_4_ only. Values represent mean ± 95% confidence intervals from four biological replicates, with 25 seeds per replicate plate. Different letters indicate significant differences among inoculum levels within each strain and day according to Tukey’s HSD test (*P* < 0.05).

For strain B18, germination percentage was significantly influenced by inoculum concentration during the early stages of germination (Day 1: F_3,12_ = 15.49, *P* < 0.001; Day 2: F_3,12_ = 8.86, *P* = 0.002). On Day 1, seeds treated with B18 reached germination levels ranging from approximately 70% to 92%, compared with approximately 60% in the MgSO_4_ control. By Day 2, germination increased to approximately 85 to 98% in inoculated treatments, whereas the control remained near 75%. At the final assessment (Day 3), all B18 treatments achieved near-complete germination (93– 100%), exceeding the control (approximately 92–94%). However, treatment differences were no longer statistically significant at this stage (F_3,12_ = 2.83, *P* = 0.083), indicating convergence toward maximum germination. Germination speed, estimated using Maguire’s Germination Index (MGI), was significantly affected by B18 inoculation (F_3,12_ = 17.21, *P* < 0.001), with the highest MGI observed at 10^7^ CFU mL^-1^. According to Tukey’s test, only the highest inoculum concentration differed significantly from the control and lower concentrations, indicating a dose-dependent enhancement of germination vigour.

In contrast, strain B14 produced a weaker but concentration-dependent stimulatory effect. Germination percentage did not differ significantly among treatments on Day 1 (F_3,12_ = 2.27, *P* = 0.133), when little or no germination was observed in either the control or most inoculated treatments. By Day 2, germination increased to approximately 24 to 38% for B14-treated seeds compared with approximately 10% for the control, with treatment effects approaching significance (F_3,12_ = 3.48, *P* = 0.050). At the final assessment (Day 3), germination reached approximately 36 to 55% across inoculum concentrations, with the highest response observed at 10^6^ CFU mL⁻¹ (approximately 55%) compared with only 22% in the control. Treatment effects were significant at this stage (F_3,12_ = 4.58, *P* = 0.023), demonstrating that B14 improved final germination despite producing a weaker response than B18. Similarly, MGI differed significantly among B14 treatments (F_3,12_ = 4.21, *P* = 0.030), with the greatest increase occurring at 10^6^ CFU mL^-1^, while the remaining inoculum concentrations produced intermediate responses.

Collectively, these results demonstrate that both endophytic *Pseudomonas* strains enhanced seed vigour relative to the MgSO_4_ control, but their effectiveness differed substantially. Strain B18 accelerated germination, increased final germination percentage, and produced the highest MGI values, indicating superior stimulation of early seed physiological processes. In contrast, B14 generated a more moderate and concentration-dependent response. The stronger performance of B18 during seed germination is consistent with its greater phosphate-solubilizing capacity and higher indole-3-acetic acid production, suggesting a stronger ability to promote early seedling establishment. To determine whether extracellular metabolites contributed to these responses, additional assays were subsequently conducted using cell-free supernatants from each strain.

### 3.5 Effects of cell-free supernatants on seed germination and radicle development

To determine whether extracellular metabolites contributed to the observed seed responses, sterile cell-free supernatants (CFS) obtained from strains B18 and B14 were evaluated in the absence of live bacterial cells. Supernatants were tested at 2.5, 5, and 10% (v/v), corresponding to 25, 50, and 100 µL mL^-1^, respectively. Water and uninoculated sterile growth medium controls were included to distinguish metabolite-mediated effects from possible nutrient carryover.

The CFS derived from strain B18 significantly affected both germination percentage (F_6,22_ = 3.68, *P* = 0.011) and radicle length (F_6,11_ = 6.42, *P* = 0.004) (Supplementary Fig. S3). Final germination increased from approximately 86% in the water control to 92% following treatment with 2.5% B18-CFS, representing the highest germination response observed among all treatments. Germination remained high at 5 and 10% CFS (approximately 90 to 91%), indicating that maximal stimulation occurred at the lowest concentration tested. Radicle growth exhibited a similar trend, increasing from approximately 5.6 cm in the control to 6.5 cm at 2.5% B18-CFS. However, radicle elongation declined at higher concentrations, reaching approximately 5.2 cm at 5% CFS and approximately 4.5 cm at 10% CFS. According to Tukey’s multiple comparison test, the 2.5% B18-CFS treatment consistently produced the strongest stimulatory response, suggesting that extracellular metabolites released by strain B18 directly promote seed germination and early root development, with maximal activity occurring at relatively low concentrations.

In contrast, the CFS derived from strain B14 also significantly influenced germination percentage (F_6,22_ = 3.70, *P* = 0.011) and radicle length (F_6,22_ = 11.26, *P* < 0.001), although the overall biological response differed markedly from that of B18. Germination percentages ranged from 63 to 75% across B14-CFS treatments, values that were comparable to or lower than the water control (77%). Radicle responses were strongly concentration dependent. While 2.5% B14-CFS moderately stimulated radicle elongation (approximately 5.6 cm), increasing the concentration substantially reduced root growth, with radicle length declining to approximately 2.6 cm at 5% CFS and being completely inhibited at 10% CFS. These findings indicate that metabolites present in B14-CFS may exert inhibitory effects at elevated concentrations.

The uninoculated medium controls did not consistently improve germination or radicle development, confirming that the positive responses observed with B18-CFS were primarily attributable to bacterial secreted metabolites rather than residual nutrients in the growth medium. Collectively, these results demonstrate that strain B18 promotes seed germination and early root growth through soluble extracellular compounds, whereas the beneficial effects associated with strain B14 are likely dependent on live-cell colonization, rhizosphere interactions, or other indirect plant–microbe processes that are not reproduced by cell-free supernatants alone.

### 3.6 Phylogenetic identification of strains B14 and B18

Maximum-likelihood analysis of partial 16S rRNA gene sequences placed both isolates within the genus *Pseudomonas* (Fig. 2). Strain B14 clustered most closely with *Pseudomonas marginalis* and *P. rhodesiae* and was further associated with related environmental species including *P. arenae* and *P. grimontii*. Bootstrap support within this clade ranged from moderate to strong (51–100%), supporting a stable phylogenetic placement of B14 among plant-associated and environmental *Pseudomonas* taxa.

**Figure 2.**
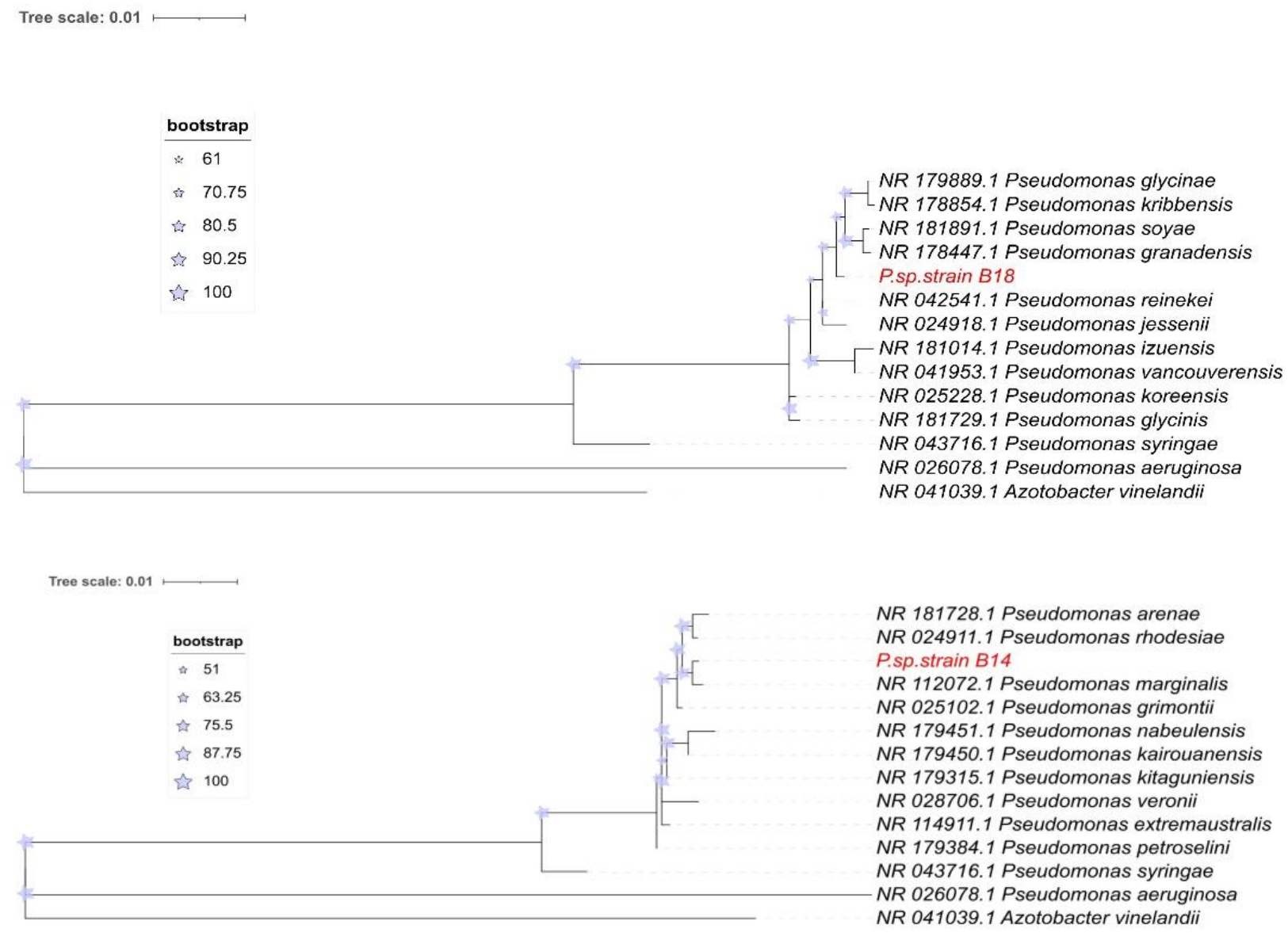
(2.1, 2.2). Maximum-likelihood phylogenetic trees based on partial 16S rRNA gene sequences showing the placement of strains B14 and B18 among closely related *Pseudomonas* reference species. Trees were inferred under the GTR+G+I model with bootstrap support indicated by node symbols. *Azotobacter vinelandii* was used as the outgroup. Strains B14 and B18 are highlighted in red.

Strain B18 formed a separate lineage associated with *Pseudomonas reinekei*, *P. jessenii*, *P. vancouverensis*, *P. uzonensis*, and closely related species. Bootstrap support within this cluster ranged from low to strong (25-100%), reflecting the close evolutionary relationships among these taxa and the limited species-level resolution provided by partial 16S rRNA gene sequences.

Neither isolate clustered with the pathogenic reference species *Pseudomonas syringae* or *P. aeruginosa*, which formed distinct branches outside the isolate-associated clades. The phylogenetic analysis therefore supports the classification of B14 and B18 as environmental or plant-associated *Pseudomonas* spp., consistent with their observed plant growth–promoting and biocontrol characteristics.

### 3.7 Greenhouse evaluation against *Verticillium dahliae*

#### 3.7.1 Plant establishment and vegetative growth

Plant emergence remained consistently high in both the absence and presence of *Verticillium dahliae*, indicating successful tuber establishment across all treatments and demonstrating that PGPR inoculation did not adversely affect early emergence (Fig. 3A).

**Figure 3.**
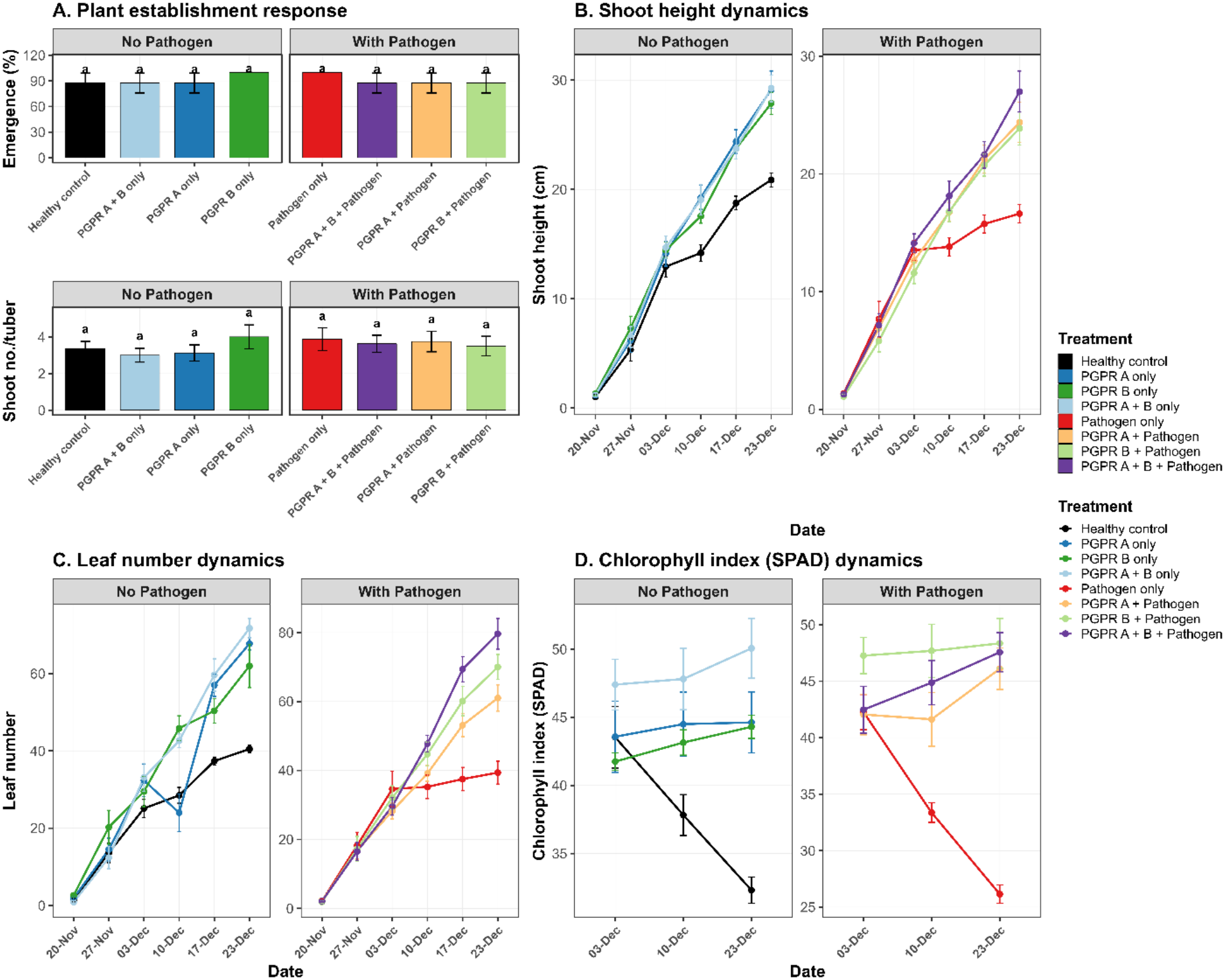
Effects of PGPR treatments on potato establishment, growth, and physiological performance under non-pathogen and pathogen-challenged conditions. (A) Plant establishment response showing emergence (%) and shoot no./tuber, (B) shoot height dynamics, (C) leaf number dynamics, and (D) chlorophyll index measured by SPAD over the experimental period. Values represent mean ± standard error (SE) from biological replicates arranged in a randomized complete block design (RCBD). Data were analyzed using RCBD ANOVA followed by Tukey’s HSD multiple comparison test. Different letters indicate significant differences among treatments within each pathogen group (*P* < 0.05).

Vegetative growth was significantly affected by treatment, particularly under pathogen challenge. Final shoot height differed significantly among treatments in both pathogen-free (F_3,21_ = 10.23, *P* < 0.001) and pathogen-challenged plants (F_3,21_ = 8.90, *P* < 0.001). In the presence of *V. dahliae*, pathogen-only plants exhibited severe growth suppression and reached only approximately 16.6 cm at the final assessment (Fig. 3B). Preventive PGPR application substantially alleviated pathogen-induced growth reduction. The combined B18 plus B14 plus pathogen treatment produced the tallest pathogen-challenged plants (27.0 cm), representing a 62.4% increase relative to the pathogen-only treatment and achieving growth levels comparable to those observed under pathogen-free conditions. Individual PGPR treatments also improved shoot growth compared with pathogen-only plants; however, the consortium consistently produced the greatest response.

Leaf production followed a similar pattern and was significantly influenced by treatment under both pathogen-free (F_3,21_ = 21.10, *P* < 0.001) and pathogen-challenged conditions (F_3,21_ = 23.33, *P* < 0.001) (Fig. 3C). Plants inoculated with *V. dahliae* alone developed the fewest leaves (39.4 leaves plant^-1^), whereas the combined B18 plus B14 plus pathogen treatment produced the highest leaf number (79.6 leaves plant⁻¹). This treatment approximately doubled leaf production relative to pathogen-only plants and exceeded the healthy untreated control by approximately 33%, demonstrating a strong recovery from pathogen-induced suppression and enhanced canopy development. The individual PGPR treatments also significantly increased leaf production compared with the pathogen-only control, although their effects were less pronounced than those of the consortium.

Chlorophyll content was strongly affected by treatment (Fig. 3D). Significant treatment effects were detected under both pathogen-free (F_3,21_ = 18.00, *P* < 0.001) and pathogen-challenged conditions (F_3,21_ = 34.57, *P* < 0.001). In the presence of *V. dahliae*, chlorophyll content declined markedly in pathogen-only plants, reaching approximately 26.2 SPAD units by the final assessment. In contrast, PGPR-treated plants maintained significantly higher chlorophyll levels throughout the experiment. The highest chlorophyll content under disease pressure was observed in the B14 plus pathogen treatment (48.4 SPAD units), representing an 84.9% increase relative to the pathogen-only control. Similarly, the B18 plus B14 plus pathogen treatment maintained elevated chlorophyll content (47.6 SPAD units), indicating that PGPR inoculation effectively reduced pathogen-associated physiological stress and preserved photosynthetic capacity.

In the absence of the pathogen, PGPR inoculation also enhanced plant performance relative to the healthy untreated control. The combined B18 plus B14 treatment produced the greatest final shoot height (∼29 cm), highest leaf number (∼70 leaves plant⁻¹), and highest chlorophyll content (∼50 SPAD units), demonstrating strong plant growth–promoting activity even under non-stress conditions.

Collectively, these results demonstrate that PGPR inoculation significantly enhanced vegetative growth and physiological performance under both healthy and pathogen-challenged conditions. The combined B18 plus B14 consortium consistently produced the strongest response across all measured parameters, indicating that the simultaneous application of functionally complementary endophytic *Pseudomonas* strains provided greater benefits than either strain alone. These findings suggest that consortium-based inoculation strategies may offer an effective approach for improving plant vigor while mitigating the detrimental effects of *V. dahliae* infection.

#### 3.7.2 Flowering and tuber production

PGPR inoculation significantly improved reproductive performance and daughter tuber production under both pathogen-free and pathogen-challenged conditions (Fig. 4). Under *Verticillium dahliae* pressure, flowering was completely suppressed in the pathogen-only treatment throughout the experiment, indicating severe impairment of reproductive development. In contrast, all PGPR-treated plants maintained flowering activity. By the final assessment, flowering incidence reached approximately 75% in the B18 plus B14 plus pathogen treatment, compared with 0% in pathogen-only plants, demonstrating substantial recovery of reproductive capacity under disease stress (Fig. 4A).

**Figure 4.**
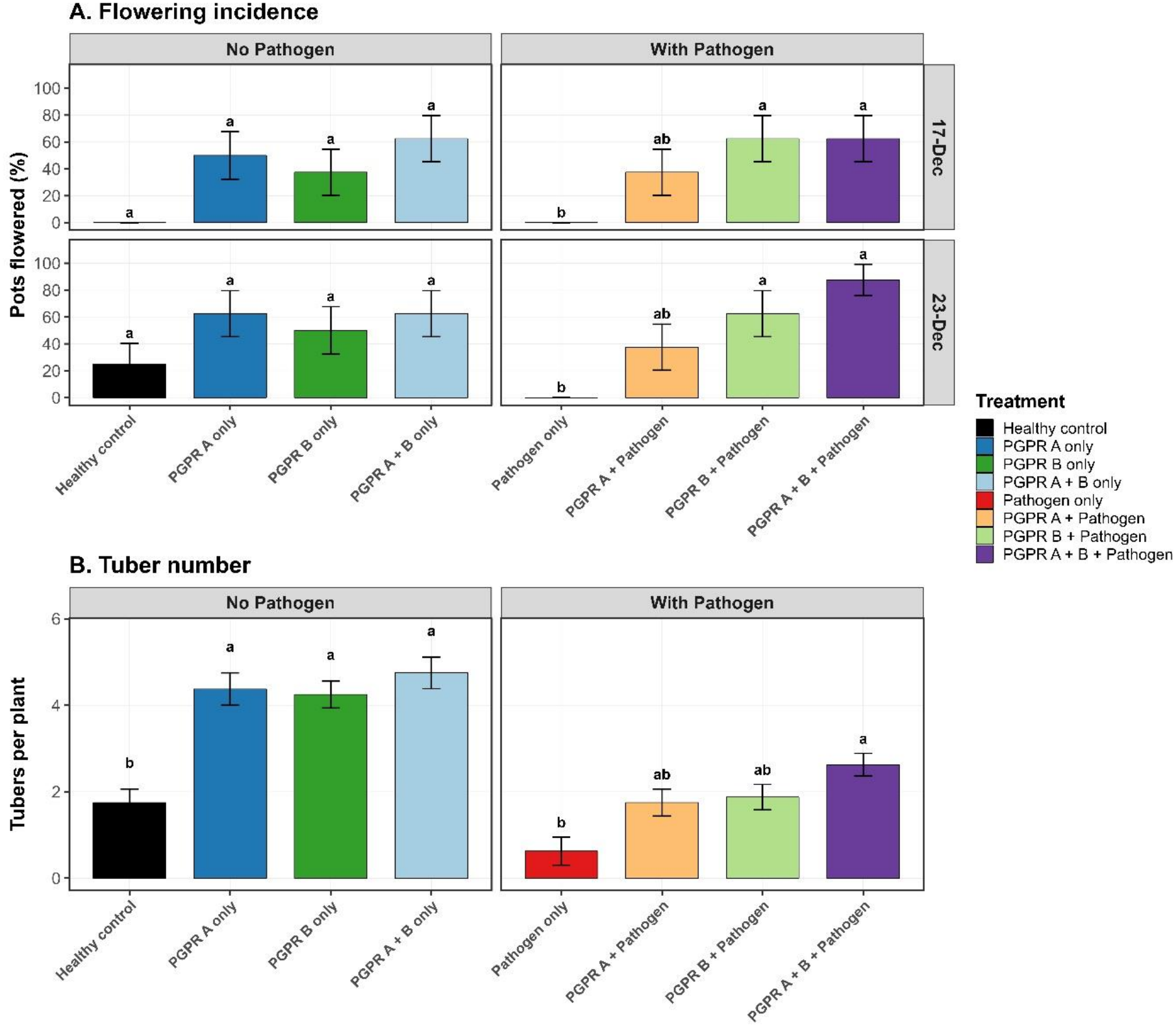
Effects of PGPR treatments on flowering and tuber production in potato. (A) Flowering incidence, calculated as the percentage of pots showing visible inflorescence formation, and (B) tuber number per plant under non-pathogen and pathogen-challenged conditions. For flowering incidence, each pot was scored as flowered or non-flowered based on the presence or absence of visible inflorescences, and the percentage of flowered pots was calculated within each treatment group. Values represent mean ± SE from biological replicates arranged in an RCBD. Data were analyzed using RCBD ANOVA followed by Tukey’s HSD test. Different letters indicate significant differences within each pathogen group (*P* < 0.05).

A similar trend was observed under pathogen-free conditions, where PGPR inoculation enhanced flowering relative to the healthy untreated control. The combined B18 plus B14 treatment produced the highest flowering incidence (∼75%), compared with ∼25% in the healthy control, indicating a positive effect of PGPR inoculation on reproductive development even in the absence of disease (Fig. 4A).

Tuber production was significantly affected by treatment under both pathogen-free (F_3,21_ = 16.06, *P* < 0.001) and pathogen-challenged conditions (F_3,21_ = 5.90, *P* = 0.004) (Fig. 4B). Under pathogen pressure, the pathogen-only treatment produced the fewest daughter tubers (0.63 tubers plant⁻¹), whereas the B18 + B14 + pathogen treatment produced the highest number (2.63 tubers plant⁻¹), representing a 4.2-fold increase relative to the pathogen-only control. Similarly, under pathogen-free conditions, the combined B18 + B14 treatment produced the greatest tuber number (4.75 tubers plant⁻¹) compared with 1.75 tubers plant⁻¹ in the healthy control, corresponding to a 171% increase (Fig. 4B).

Overall, PGPR inoculation mitigated disease-associated reductions in flowering and tuberization while simultaneously enhancing reproductive performance under healthy conditions. The B18 + B14 consortium consistently produced the strongest response, highlighting the benefits of combining functionally complementary endophytic *Pseudomonas* strains.

#### 3.7.3 Biomass accumulation and root system architecture

PGPR inoculation significantly enhanced biomass accumulation and root development under both pathogen-free and *Verticillium dahliae*–challenged conditions (Fig. 5). Under pathogen pressure, total fresh biomass differed significantly among treatments (F_3,17_= 4.55, *P* = 0.016). The pathogen-only treatment produced the lowest biomass (11.4 g plant⁻¹), whereas the B18 + B14 + pathogen treatment produced the highest value (16.0 g plant⁻¹), representing a 40.0% increase relative to the pathogen-only control (Fig. 5A). Total dry biomass was similarly affected by treatment (F_3,16_ = 3.45, *P* = 0.042), increasing from 1.64 g plant⁻¹ in pathogen-only plants to 2.45 g plant⁻¹ in consortium-treated plants, corresponding to a 49.0% improvement (Fig.5B).

**Figure 5.**
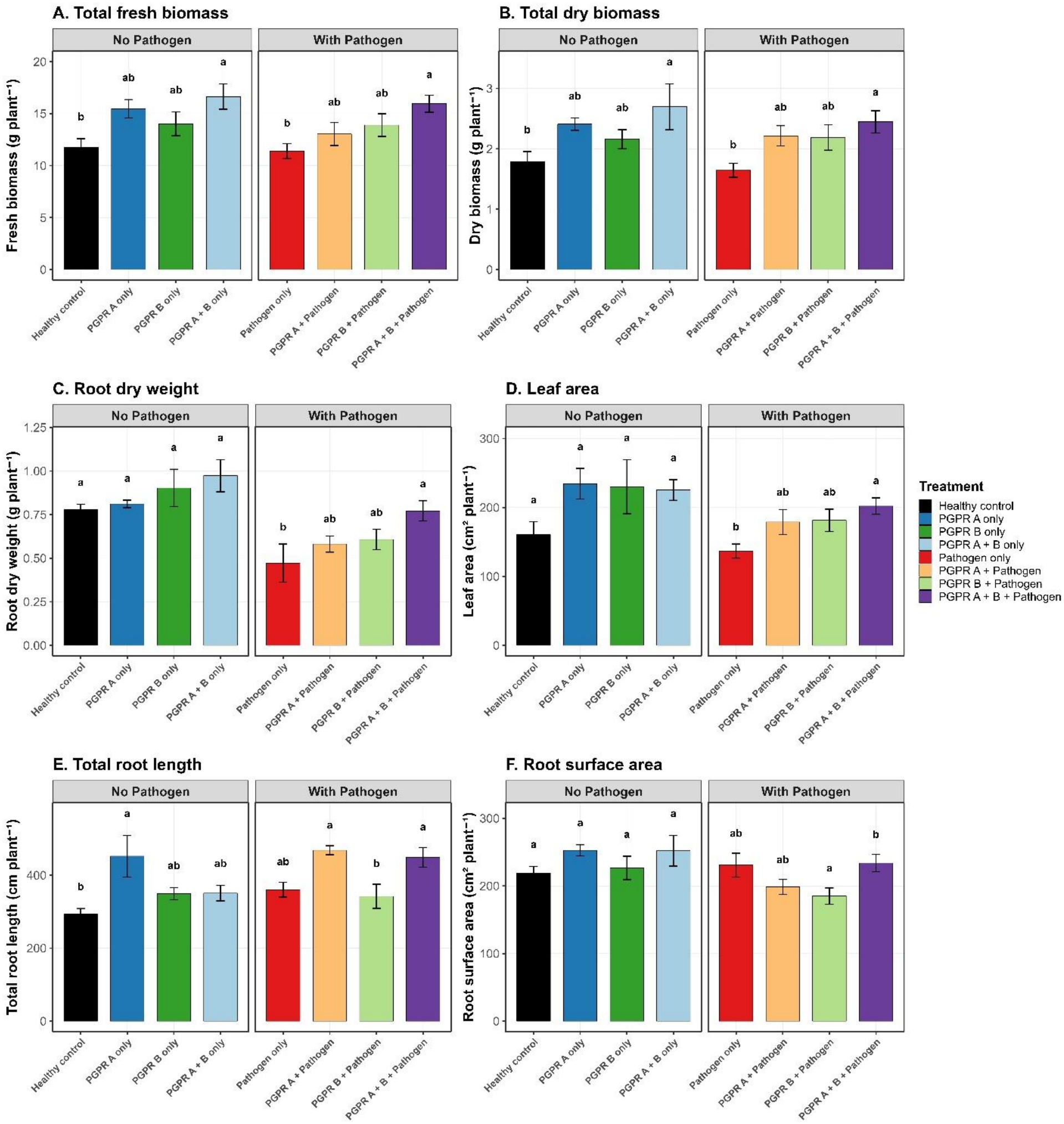
Effects of endophytic *Pseudomonas* strains on potato biomass accumulation and root system architecture under non-pathogen and *Verticillium dahliae*–challenged conditions. (A) Total fresh biomass, (B) total dry biomass, (C) root dry weight, (D) leaf area, (E) total root length, and (F) root surface area measured at final harvest. Values represent mean ± standard error (SE) from eight biological replicates arranged in a randomized complete block design (RCBD). Treatments included Healthy control, PGPR A (*Pseudomonas* strain B18), PGPR B (*Pseudomonas* strain B14), PGPR A + B consortium, Pathogen only (*V. dahliae*), PGPR A + Pathogen, PGPR B + Pathogen, and PGPR A + B + Pathogen. Data were analyzed using RCBD ANOVA with treatment as a fixed effect and block as a blocking factor, followed by Tukey’s honestly significant difference (HSD) test (*P* < 0.05). Different letters indicate significant differences among treatments within each pathogen group.

Leaf area was significantly influenced by treatment under pathogen challenge (F_3,17_ = 3.79, *P* = 0.030). The pathogen-only treatment exhibited the smallest leaf area (136.7 cm^2^ plant^-1^), whereas the B18 plus B14 plus pathogen treatment reached approximately 202.0 cm^2^ plant^-1^, representing a 47.8% increase (Fig. 5D). Root dry weight also increased substantially in response to PGPR inoculation and approached statistical significance under pathogen pressure (F_3,17_ = 3.12, *P* = 0.053). Consortium-treated (B18 plus B14) plants produced the greatest root biomass (0.77 g plant^-1^), representing a 63.3% increase relative to pathogen-only plants (Fig. 5C).

Root architectural traits showed similar responses. Total root length differed significantly among treatments under both pathogen-free (F_3,15_ = 3.96, *P* = 0.029) and pathogen-challenged conditions (F_3,17_ = 5.19, *P* = 0.010). Under disease pressure, consortium-treated plants developed more extensive root systems, reaching approximately 449 cm plant^-1^ compared with 360 cm plant⁻¹ in pathogen-only plants (Fig. 5E). Root surface area was also significantly affected under pathogen challenge (F_3,17_ = 3.92, *P* = 0.027), indicating improved root system development despite disease pressure (Fig. 5F).

In the absence of the pathogen, PGPR treatments also promoted plant growth relative to the healthy untreated control. The B18 and B14 consortium produced the greatest total fresh biomass (∼16.8 g plant^-1^), total dry biomass (∼2.7 g plant^-1^), leaf area (∼225 cm^2^ plant^-1^), and root biomass (∼0.97 g plant^-1^), demonstrating strong plant growth–promoting activity even under non-stress conditions (Fig. 5A, 5B, 5C).

Among the individual strains, B18 generally produced stronger effects on root development than B14, particularly for total root length and root surface area. Nevertheless, the combined B18 and B14 treatment consistently generated the greatest and most balanced improvements across both aboveground and belowground traits. Similar trends were observed for leaf fresh weight, stem fresh weight, leaf dry weight, stem dry weight, root volume, and average root diameter (Supplementary Fig. S4). Collectively, these results demonstrate that endophytic *Pseudomonas* strains B18 and B14 substantially enhanced biomass accumulation and root system architecture while mitigating the detrimental effects of *V. dahliae* on potato growth.

#### 3.7.6 Disease suppression by PGPR treatments

Disease progression, expressed as the area under the disease progress curve (AUDPC), was significantly affected by PGPR treatment under *Verticillium dahliae* challenge (F_3,21_ = 14.59, *P* < 0.001) (Fig. 6). The pathogen-only treatment exhibited the highest cumulative disease severity, with a mean AUDPC value of 40.94, indicating severe wilt development in the absence of biological protection.

**Figure 6.**
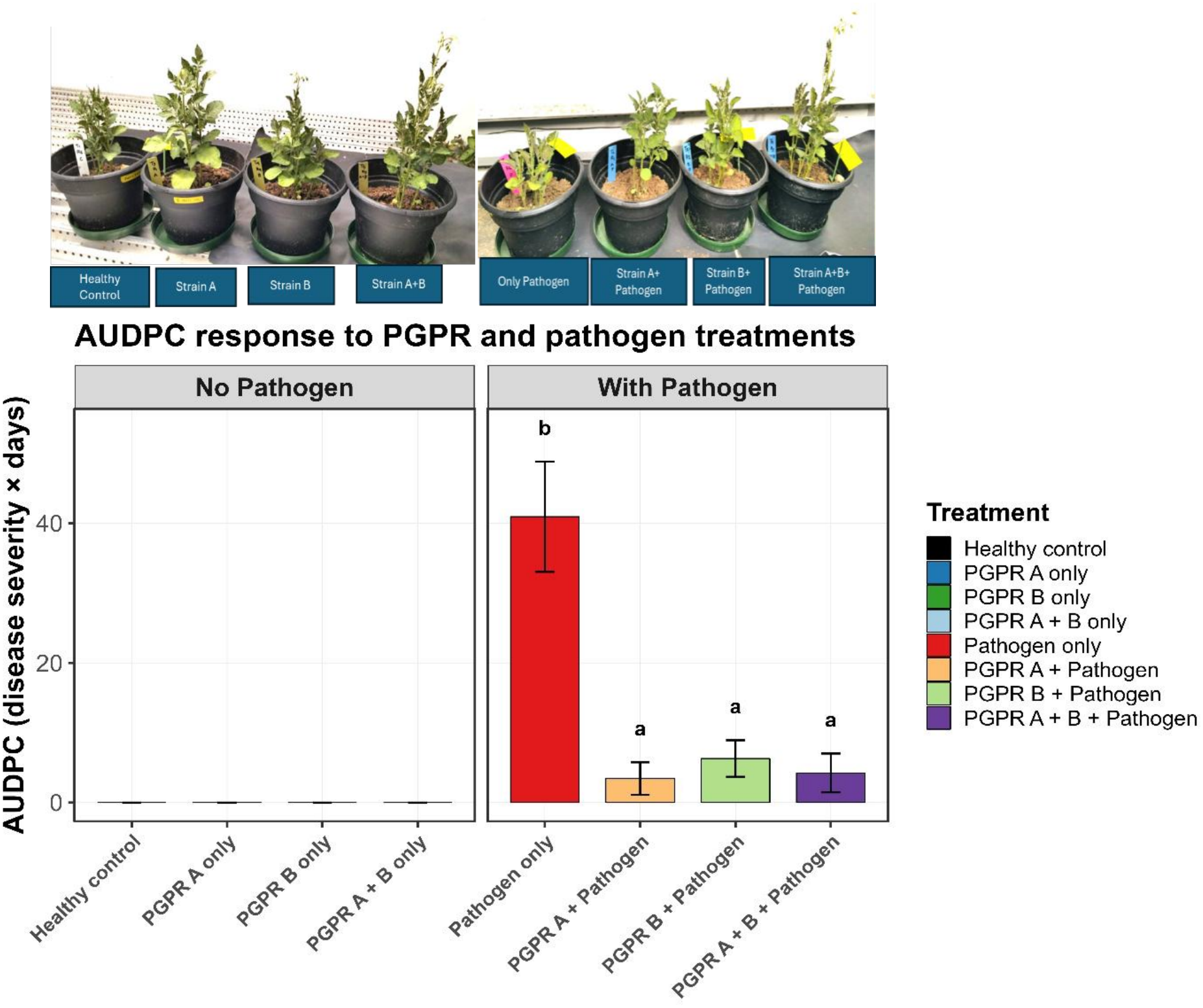
Effects of PGPR treatments on *Verticillium* wilt progression in potato. Disease progression was summarized as the area under the disease progress curve (AUDPC) under non-pathogen and *Verticillium dahliae*– challenged conditions. Values represent mean ± SE from biological replicates arranged in an RCBD. Data were analyzed using RCBD ANOVA followed by Tukey’s HSD test. Different letters indicate significant differences within the pathogen-challenged group (*P* < 0.05). No statistical letters are shown for the non-pathogen group because AUDPC values were zero across treatments.

Preventive PGPR inoculation substantially reduced disease progression relative to the pathogen-only control. AUDPC values were 3.44, 6.31, and 4.25 for the B18 plus pathogen, B14 plus pathogen, and B18 plus B14 plus pathogen treatments, respectively. These values corresponded to reductions in cumulative disease severity of 91.6, 84.6, and 89.6% relative to the pathogen-only treatment. Although the B18 plus pathogen treatment produced the numerically lowest AUDPC value, Tukey’s multiple comparison test detected no significant differences among the three PGPR-protected treatments (*P* > 0.05), indicating that all inoculation strategies provided similarly effective suppression of Verticillium wilt.

No disease symptoms were observed in the non-inoculated treatments, and AUDPC values remained at or near zero throughout the experiment. Consequently, statistical analysis was not applicable for the non-pathogen treatments because disease severity did not develop under these conditions.

Overall, the results demonstrate that both individual strains and their combined application provided strong and consistent biological control of *V. dahliae*, reducing cumulative disease severity by more than 84% compared with the pathogen-only control. While disease suppression was statistically comparable among PGPR treatments, the B18 plus B14 consortium consistently produced superior plant growth, biomass accumulation, root development, and tuber production throughout the experiment. These findings suggest that the principal advantage of the consortium lies not in greater disease suppression alone, but in its ability to simultaneously support plant growth and maintain crop performance under pathogen pressure.

### 3.8 Integrated performance of single-strain and consortium treatments

Across laboratory, growth chamber, and greenhouse experiments, the two endophytic *Pseudomonas* strains exhibited distinct but complementary functional profiles. Strain B18 consistently demonstrated stronger plant growth–promoting characteristics, including greater phosphate solubilization, higher indole-3-acetic acid production, stronger seed germination stimulation, improved shoot and root growth, and enhanced physiological performance. In contrast, strain B14 displayed stronger antagonistic activity against *Verticillium* spp. *in vitro* and contributed substantially to the maintenance of chlorophyll content and plant health under pathogen challenge.

Although both strains individually improved plant performance and reduced disease severity, their combined application consistently produced the most balanced and comprehensive response across measured agronomic traits. The B18 plus B14 consortium enhanced shoot growth, leaf production, biomass accumulation, root development, flowering, tuber production, and disease tolerance under both healthy and pathogen-challenged conditions. Importantly, while all PGPR treatments provided statistically comparable suppression of Verticillium wilt, the consortium consistently generated superior overall plant performance, indicating benefits beyond disease control alone.

Collectively, these findings support the concept of functional complementarity, whereby strains possessing different biological capabilities contribute distinct but synergistic benefits to plant performance. The results further suggest that carefully selected multi-strain PGPR consortia may provide a more robust strategy for improving crop productivity and resilience than single-strain inoculants. Consequently, the B18 plus B14 consortium represents a promising candidate for the integrated management of Verticillium wilt and the development of sustainable potato production systems.

## 4. Discussion

Our study shows that stress-adapted endophytic *Pseudomonas* strains isolated from unmanaged wild lupin roots have considerable potential to promote potato growth and suppress Verticillium wilt under glasshouse conditions. Both isolates exhibited multiple plant growth–promoting (PGP) traits, inhibited *Verticillium* species *in vitro*, enhanced plant performance, and reduced disease severity. The two strains displayed distinct functional profiles: B18 showed stronger growth-promoting characteristics, while B14 exhibited greater direct antifungal activity. Notably, the combined application of both strains consistently led to the greatest improvements in plant growth, physiological performance, biomass accumulation, tuber production, and disease suppression. These data suggest that functionally complementary microbial inoculants may deliver broader benefits than single-strain applications.

The antagonistic activity observed against *V. dahliae, V. longisporum*, and *V. albo-atrum* is consistent with previous reports identifying *Pseudomonas* species as effective biological control agents against soil-borne fungal pathogens (Haas and Défago, 2005; Weller, 2007). Of the two isolates, B14 consistently manifested greater inhibition of fungal growth in dual-culture assays, indicating stronger direct antagonistic potential. *Pseudomonas* species are recognized for producing a range of antimicrobial metabolites, volatile compounds, siderophores, and extracellular enzymes that may contribute to fungal suppression (Mercado-Blanco and Bakker, 2007; Pieterse et al., 2014). However, as antimicrobial compounds were not directly characterized in this study, the particular mechanisms underlying fungal inhibition remain undetermined. Consequently, the noted suppression could be regarded as evidence of biocontrol potential rather than confirmation of specific antimicrobial pathways.

Beyond pathogen suppression, both isolates exhibited multiple plant-beneficial functions, including phosphate solubilization, indole-3-acetic acid (IAA) production, nitrogen fixation-associated activity, siderophore production, and hydrogen cyanide production. These traits are typically linked to greater nutrient acquisition, root development, and plant vigour (Lugtenberg and Kamilova, 2009; Glick, 2012). Strain B18 consistently surpassed B14 in phosphate solubilization, IAA production, siderophore activity, and promotion of seed germination, showing a stronger role in supporting plant growth and early seedling establishment. The observed improvements in shoot growth, chlorophyll content, biomass accumulation, and root development in greenhouse experiments are consistent with previous studies that have demonstrated the positive effects of multifunctional *Pseudomonas* strains on plant growth (Weller, 2007; Mercado-Blanco and Bakker, 2007). However, since nutrient uptake, phytohormone concentrations, and plant physiological pathways were not directly measured, these mechanisms should be considered plausible contributors rather than confirmed causes of the observed responses.

An important finding of this study is the better performance of the B18 plus B14 consortium. While each strain improved plant performance and reduced disease severity, the combined treatment consistently produced the strongest responses across nearly all measured parameters. This result indicates that the consortium’s enhanced performance results from functional complementarity between the two isolates. The strain B18 exhibited stronger plant growth–promoting activity, whereas B14 demonstrated greater antifungal capacity. Their combined application appears to provide a wider range of beneficial functions than either strain alone. Similar results have been reported for microbial consortia (Tao et al., 2020), where microorganisms with distinct ecological functions outperform individual strains by simultaneously aiding nutrient acquisition, plant growth promotion, pathogen suppression, and stress mitigation (Bashan et al., 2014; Niu et al., 2017; Berg et al., 2021).

The concept of functional complementarity has received increasing attention in microbiome-based agriculture. Instead of relying on a single primary mechanism, effective microbial consortia may operate through the division of environmental roles, niche differentiation, and the integration of multiple beneficial activities within the rhizosphere (Trivedi et al., 2020). Although direct interactions between B18 and B14 were not investigated, the consistent superiority of the consortium treatment under both healthy and pathogen-challenged conditions indicates that these isolates occupy complementary functional niches. Such combinations may provide greater stability and broader efficacy than single-strain inoculants, especially under shifting environmental conditions. Therefore, the development of carefully selected microbial consortia may offer a promising strategy to enhance both biological control and crop productivity.

An additional significant observation was the retention of biological activity in the cell-free supernatant (CFS) of strain B18. Sterile CFS considerably enhanced seed germination and radicle elongation in the absence of live bacterial cells, indicating that extracellular metabolites contributed to the observed plant responses. Comparable effects have been reported for bacterial signaling molecules, phytohormones, volatile organic compounds, lipopeptides, and other low-molecular-weight metabolites produced by plant-associated microorganisms (Lee et al., 2009; Subramanian and Smith, 2015; Raaijmakers and Mazzola, 2012). Although the specific active compounds responsible for the stimulatory effects remain unidentified, these results suggest that strain B18 may be a promising source of biologically active metabolites with agricultural potential.

The capacity of sterile bacterial culture filtrates to reproduce beneficial plant responses has significant practical consequences. Metabolite-based biostimulants may offer several advantages over live microbial inoculants, including improved formulation stability, extended shelf life, simplified storage requirements, and reduced variability associated with microbial establishment under field conditions (Backer et al., 2018). Subsequent research should therefore prioritize bioassay-guided fractionation of B18 cell-free supernatants, combined with metabolomic and chemical characterization, to identify the compounds accountable for growth promotion. These studies could facilitate the advancement of next-generation biological products and improve understanding of the molecular basis of plant–microbe interactions.

The origin of both isolates from unmanaged wild lupin plant populations that survive in cold, environmentally challenging conditions may be relevant to their performance. Wild plants frequently harbour microbiomes enriched in traits associated with nutrient acquisition, environmental acclimation, and pathogen suppression (Compant et al., 2010; Hardoim et al., 2015). Those environments may serve as natural reservoirs for microorganisms with traits advantageous for agricultural applications. The successful performance of B14 and B18 supports the increasing recognition that stress-adapted natural ecosystems are valuable sources of elite microbial inoculants for sustainable agriculture.

In summary, the findings show that stress-adapted endophytic *Pseudomonas* strains isolated from unmanaged wild lupin roots can significantly improve potato growth and reduce Verticillium wilt severity under growth-chamber conditions. The better performance of the B18 plus B14 consortium demonstrates the importance of combining strains with complementary functional traits and supports the wider concept that multifunctional microbial communities may offer more consistent agricultural benefits than individual isolates. These results add to the expanding body of evidence supporting microbiome-based strategies for sustainable disease management and crop production.

Some specific considerations should be taken into account when interpreting the results of this study. First, all experiments were carried out under controlled environmental conditions; therefore, field validation is necessary before any kind of agronomic recommendations can be made. Environmental variation, soil characteristics, indigenous microbial communities, and agricultural management practices may influence inoculant performance under commercial production conditions. Second, taxonomic identification was based on partial 16S rRNA gene sequencing, which provides limited species-level resolution within the genus *Pseudomonas*. Whole-genome sequencing would improve taxonomic resolution and facilitate the identification of genes associated with plant growth promotion, secondary metabolite biosynthesis, biocontrol activity, habitat adaptation, and biosafety.

Third, although multiple plant growth–promoting traits and strong antagonistic activity were observed, the molecular and biochemical mechanisms underlying these responses were not directly investigated. Future research needs to be conducted to integrate whole-genome sequencing, in combination of plant metabolomic profiling, transcriptomics, and rhizosphere colonization analyses to identify the genes, metabolites, and pathways accounting for disease suppression and plant growth promotion. Specifically, characterization of the bioactive compounds present in B18 cell-free supernatants may detect novel metabolites suitable for development as microbial biostimulants. Additionally, long-term persistence studies and investigations into interactions with native microbial communities will be key to assessing the ecological stability, effectiveness, and feasibility of deploying these strains under field conditions.

## 5. Acknowledgments

The authors gratefully acknowledge the financial support provided by McCain Foods and the support of the Eastern Canada Oilseeds Development Alliance (ECODA) project. We thank all members of the Donald Smith’s Laboratory, Department of Plant Science, McGill University, for their assistance with greenhouse experiments, laboratory work, technical support, and valuable scientific discussions throughout this study. The authors are especially grateful to Professor Donald L. Smith for his guidance, mentorship, and continued support during the project.

## Supplementary data

**Supplementary Figure S1.**
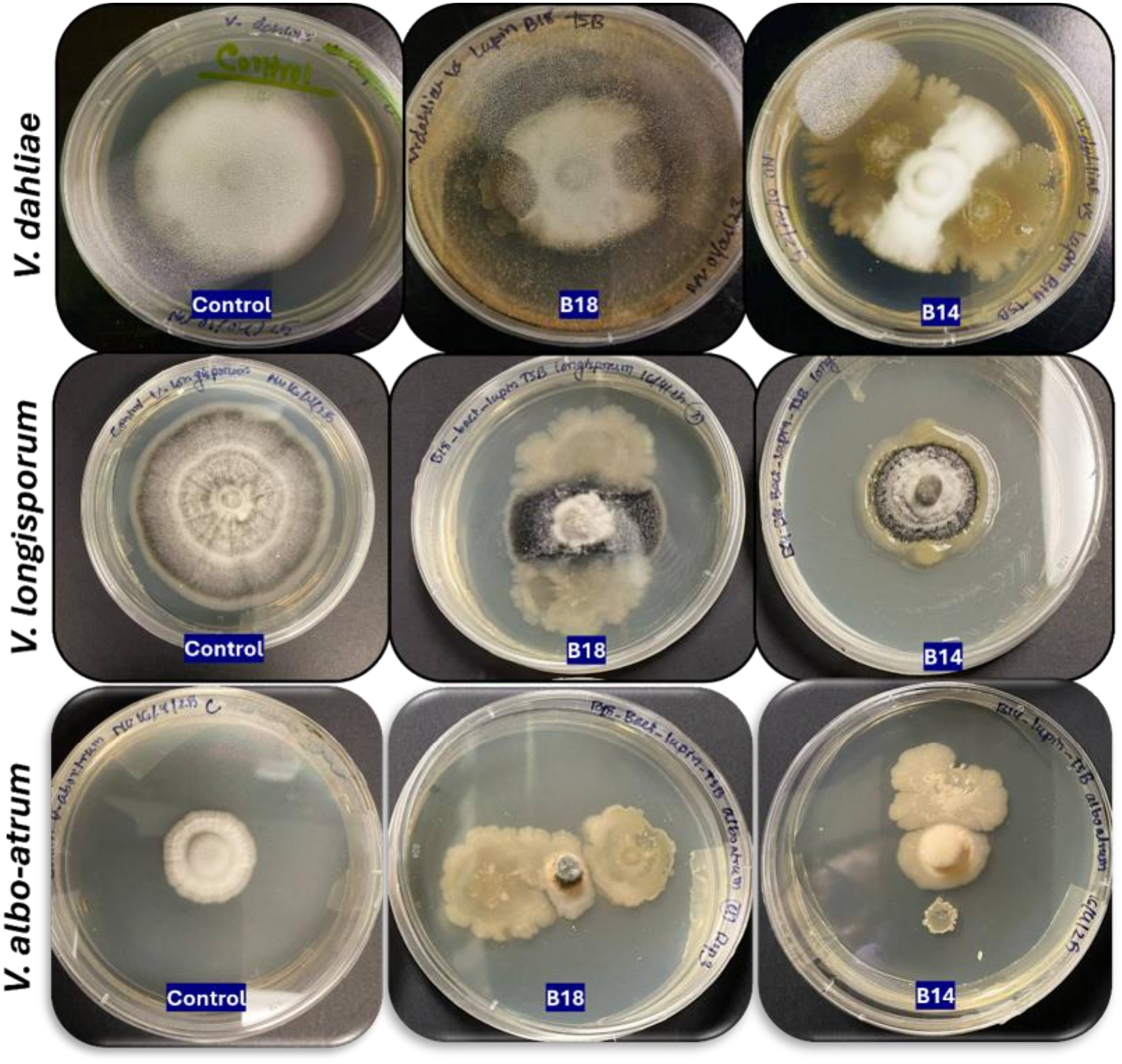
Representative dual-culture antagonism assays of isolates B18 and B14 against three *Verticillium* species after 4 weeks of incubation. Colony diameter used for quantitative inhibition analysis was measured after 2 weeks of incubation, whereas the images shown here were captured after 4 weeks to better illustrate sustained fungal growth suppression, altered colony morphology, and long-term interaction patterns. Top row: *Verticillium dahliae*; middle row: *V. longisporum*; bottom row: *V. albo-atrum*. In each row, the left plate represents the untreated fungal control, while the center and right plates show co-culture with isolates B18 and B14, respectively.

**Supplementary Figure S2.**
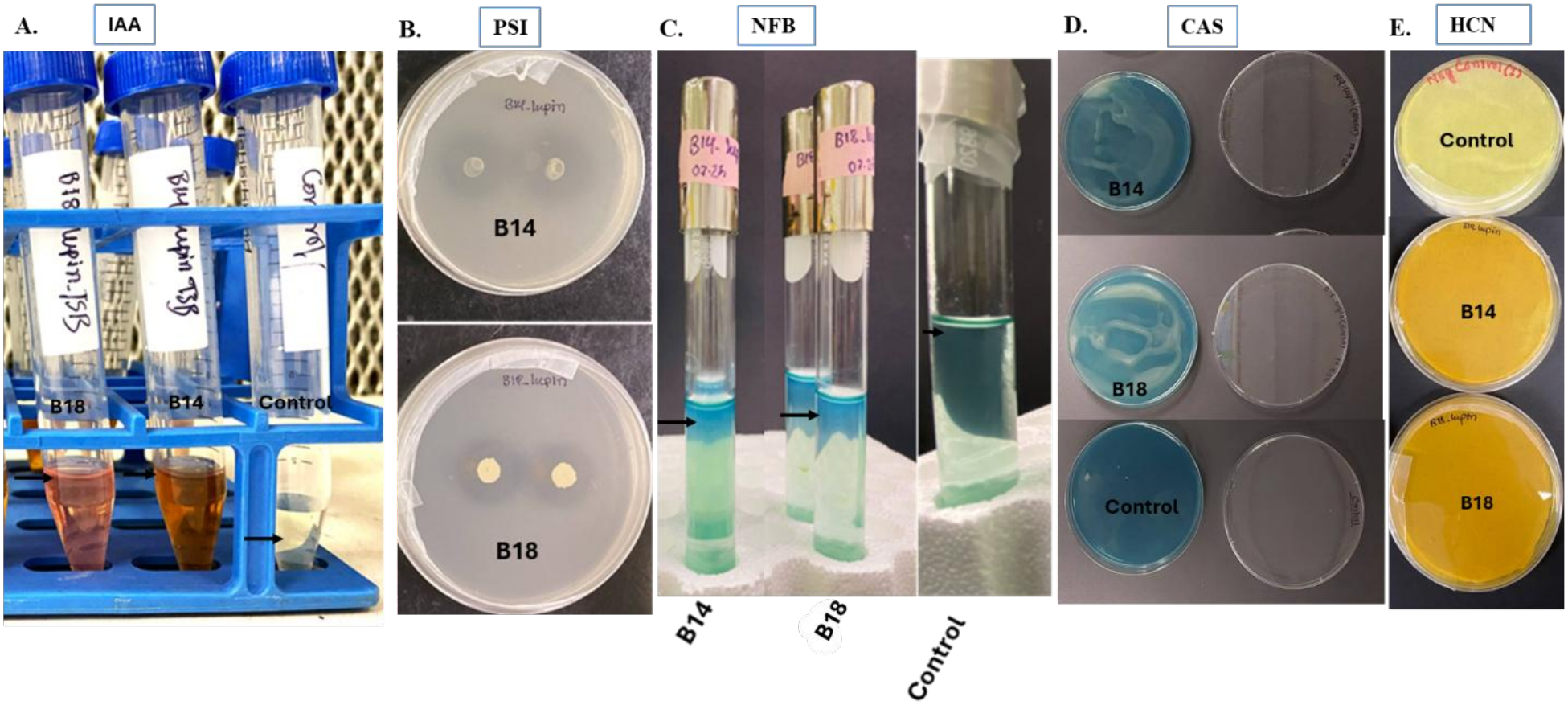
Phenotypic and biochemical characterization of endophytic *Pseudomonas* isolates B14 and B18 used in this study. Representative images showing preliminary plant growth–promoting and functional trait screening of the selected isolates. From left to right: (A) indole-3-acetic acid (IAA) production in broth culture indicated by development of pink to reddish coloration after Salkowski reagent reaction; (B) phosphate solubilization on Pikovskaya’s agar shown by halo formation surrounding colonies; (C) qualitative nitrogen fixation-associated growth in semi-solid NFb medium indicated by pellicle formation near the oxygen gradient; (D) siderophore production on chrome azurol S (CAS) medium indicated by color change around colonies; and (E) colony pigmentation and growth morphology on agar medium. Sterile uninoculated controls were included for comparison. These assays were used as preliminary screening tools to identify isolates with multiple plant growth–promoting and biocontrol-associated traits prior to greenhouse evaluation.

**Supplementary Figure S3.**
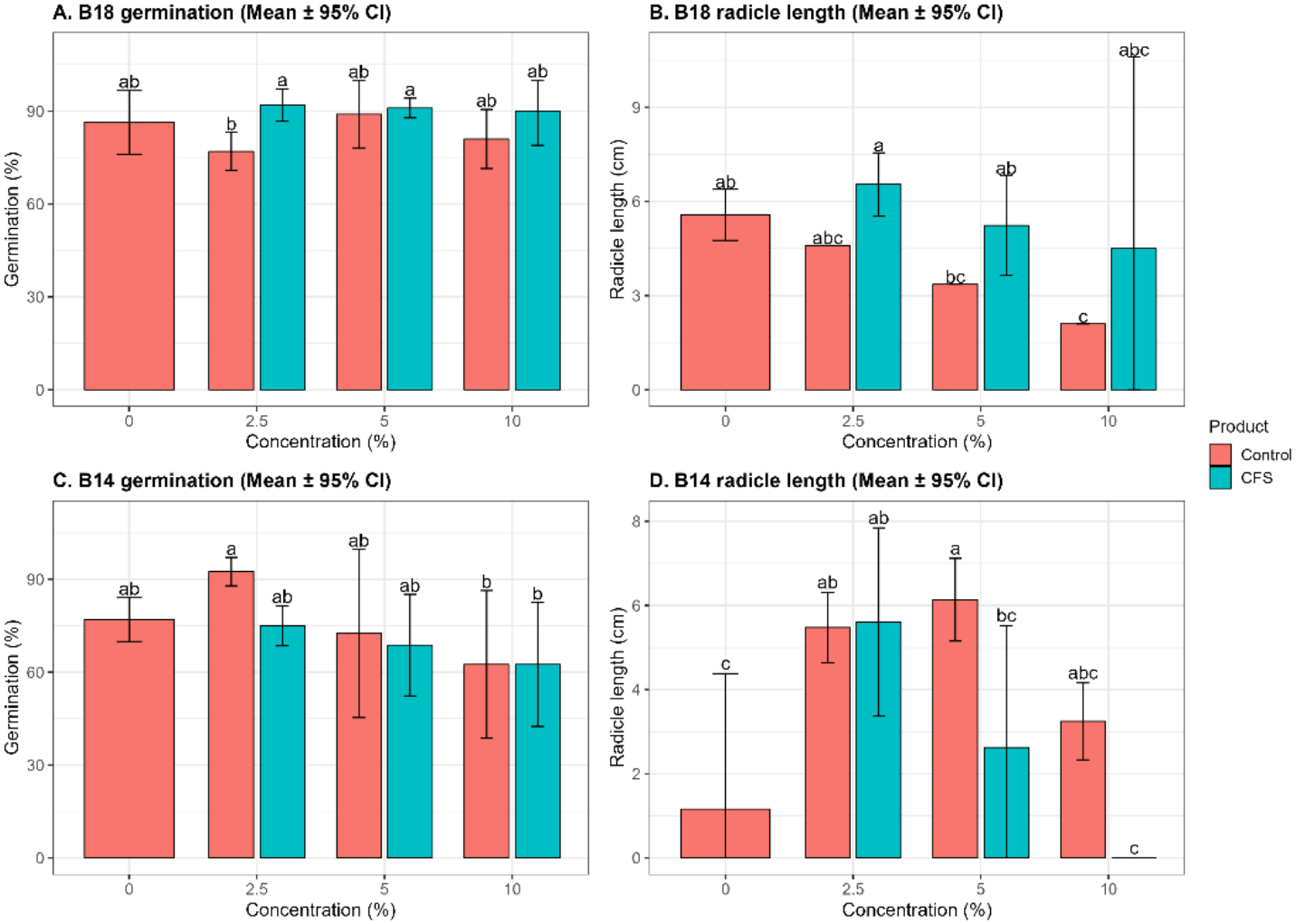
Effects of cell-free supernatant (CFS) from bacterial isolates B18 and B14 on seed germination and radicle growth at different concentrations. Four independent biological replicates were used for each treatment, with 25 seeds per replicate plate. Bars represent mean ± 95% confidence interval (CI). Different lowercase letters above bars indicate statistically significant differences among treatments within each panel according to one-way ANOVA followed by Tukey’s HSD post hoc test (*P* < 0.05). (A) Germination percentage of seeds treated with CFS from isolate B18 at 0, 2.5, 5, and 10% concentrations compared with the untreated control. (B) Radicle length of germinated seeds treated with B18 CFS at corresponding concentrations. (C) Germination percentage of seeds treated with CFS from isolate B14 at 0, 2.5, 5, and 10% concentrations compared with the untreated control. (D) Radicle length of germinated seeds treated with B14 CFS at corresponding concentrations. Overall, B18 CFS showed positive effects on germination and radicle elongation at moderate concentrations, whereas higher concentrations of B14 CFS reduced radicle growth, demonstrating concentration-dependent responses of seedling development to bacterial extracellular metabolites.

**Figure S4.**
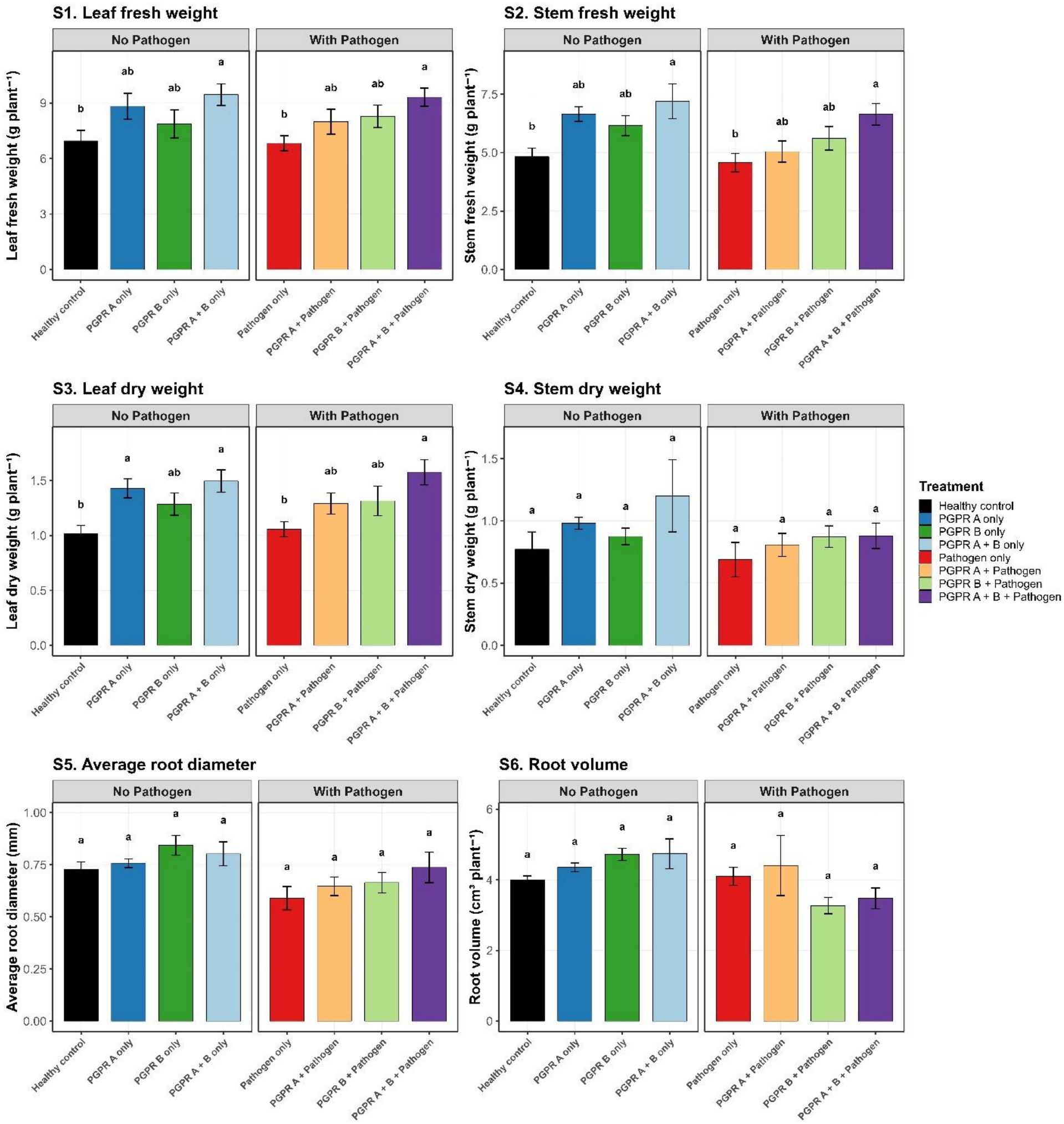
Effects of endophytic *Pseudomonas* strains on additional biomass and root architecture traits of potato under non-pathogen and *Verticillium dahliae*–challenged conditions. Panels show leaf fresh weight, stem fresh weight, leaf dry weight, stem dry weight, average root diameter, and root volume measured at final harvest. Values represent mean ± SE from eight biological replicates arranged in a randomized complete block design (RCBD). Data were analyzed using RCBD ANOVA followed by Tukey’s HSD test (*P* < 0.05). Different letters indicate significant differences among treatments within each pathogen group.

